# Increases in canopy mortality and their impact on the demographic structure of Europe’s forests

**DOI:** 10.1101/2020.03.30.015818

**Authors:** Cornelius Senf, Julius Sebald, Rupert Seidl

## Abstract

Pulses of tree mortality have been reported for many ecosystems across the globe recently. Yet, large-scale trends in tree mortality remain poorly quantified. Manually analyzing more than 680,000 satellite image chips at 19,896 plot locations, we here show that forest canopy mortality in Europe has continuously increased since 1985 (+1.5 ± 0.28 % yr^-1^), with the highest canopy mortality rate of the past 34 years observed in 2018 (1.14 ± 0.16 %). Using simulations, we demonstrate that a continued increase in canopy mortality will strongly alter forest demography, with the median forest age falling below 30 years in more than 50% of Europe’s countries by 2050. These demographic changes can have substantial cascading effects on forest regeneration, biodiversity, and carbon storage. The current trend of increasing canopy mortality is thus challenging the future of Europe’s forests, and should be a key priority of forest policy and management.

## Introduction

Tree mortality is a key demographic process in forest ecosystems. It drives natural ecosystem dynamics and occurs at different spatial scales, from the level of individual trees (Franklin *et al*. 2002; Lutz and Halpern 2006) to large-scale pulses of mortality, referred to as disturbances (Turner 2010). While many causes of tree death are natural (e.g., resource limitation, wildfire or outbreaks of native insects), there are also human causes of tree death related to land-use, e.g., societal demand for biomass (Curtis *et al*. 2018) or the introduction of alien forest pests (Roy *et al*. 2014). Both natural and human causes of tree death have been found to increase across many forest ecosystem globally (van Mantgem *et al*. 2009; Allen *et al*. 2010; Seidl *et al*. 2014; Roy *et al*. 2014), raising concerns about the potential impacts on the health and resilience of global forests (Trumbore *et al*. 2015; Johnstone *et al*. 2016).

Large-scale changes in tree mortality can have substantial and widespread impacts on forest ecosystems. Trees are long-lived and regenerate slowly, and an increase in tree mortality can shift the population structure of forests towards younger trees (Anderson-Teixeira *et al*. 2013;). Globally, the share of young forests has already increased from 11.3% to 33.6% since 1900 (McDowell et al. 2020). Such shifts in forest demography can have ripple effects on forest functions and services (Anderegg *et al*. 2012; Thom and Seidl 2016). For example, old forests represent important habitat for many forest-dwelling species (Bengtsson *et al*. 2000), and homogenizing forests towards a younger population structure will therefore have negative impacts on overall forest biodiversity. Old forests are likewise important for carbon storage (Zhou *et al*. 2006; Luyssaert *et al*. 2008), and a shift in forest demography could have detrimental impacts on the carbon storage potential of forests. Given the substantial and widespread impacts of increasing tree mortality on global forests ecosystems, it is essential to monitor and understand changes in tree mortality from the individual to the continental scale.

For Europe there is accumulating evidence that tree mortality is increasing (Schelhaas *et al*. 2003; Seidl *et al*. 2014; Senf and Seidl 2018; Senf *et al*. 2018). This increase in tree mortality might be explained by an increased utilization of Europe’s forest resources (Forest Europe 2015; Senf *et al*. 2018) but also by more frequent and severe natural disturbances (Schelhaas *et al*. 2003; Seidl *et al*. 2014). Both the increased utilization of Europe’s forest resources and increasing natural disturbances can potentially impact forest demography and thus the functioning and ecosystem services of Europe’s forests (Nabuurs *et al*. 2013; Pedroli *et al*. 2013; Seidl *et al*. 2014; Searchinger *et al*. 2018). However, most of the evidence for increasing tree mortality in Europe relies on compilations of grey literature (Schelhaas *et al*. 2003; Seidl *et al*. 2014) or focusses on regional trends (Senf and Seidl 2018; Senf *et al*. 2018). It thus remains unclear if and how tree mortality has changed across continental Europe, and how those changes might impact Europe’s forest demography. We here address these knowledge gaps by (1) manually interpreting 680,000 satellite image chips at 19,896 plot locations covering 210 Mill. ha of forest area and a 34-year period from 1985 to 2018, in order to robustly quantify rates and trends in forest canopy mortality for continental Europe; and by (2) applying simulations to determine how trends in forest canopy mortality could affect the demography of Europe’s forests under scenarios of both stabilizing and increasing canopy mortality.

## Materials and Methods

### Estimating canopy mortality rates

We used a stratified random sampling design to select plots for satellite image interpretation across continental Europe. Continental Europe here includes 35 countries (Table S1 and Figure S1) with a minimum land area of 10,000 km^2^ (i.e., excluding Lichtenstein, Luxembourg, Monte Carlo and Malta). A plot was defined as a 30 × 30 m square corresponding to a Landsat satellite pixel. We used an equalized stratified sampling design, randomly placing 500 plots within forest areas of each country. We used an equalized stratified sampling design over a proportional sampling design because sampling proportional to forest area would have led to sample sizes too large for manual interpretation in some countries (e.g., Finland, Sweden), whereas only few samples would have been placed into others (e.g., Denmark, the Netherlands). Forest areas were determined using an existing Landsat-based forest cover map that identifies all areas which have been forested at some point in time during the study period 1985 to 2018. Using a forest cover map for stratifying our sampling greatly improved sampling efficiency, but also led to selecting plots falsely identified as being forested. We excluded these plots during satellite image interpretation, resulting in varying realized sample sizes per country (Table S1). Data for six countries (Austria, Czechia, Germany, Poland, Slovakia and Switzerland) were taken from a previous study (Senf *et al*. 2018), which used a similar sampling design but larger sample sizes. To avoid loss of information from down-sampling, we used the full sample sizes for those countries, but tested whether our results remained consistent across the pan-European dataset (see Figure S13).

For each plot we manually interpreted temporal-spectral profiles of all satellite imagery available in the Landsat archive to assess whether a canopy mortality event occurred at this specific plot location at any given year of the study period. The approach follows image processing routines and image interpretation protocols developed in a previous study (Senf *et al*. 2018) and we here only give the salient details needed for understanding our approach. A canopy mortality event was defined as any loss of canopy (e.g., biotic natural disturbance, abiotic natural disturbance, regular timber harvest, sanitation logging) that resulted in an identifiable change in the canopy’s spectral reflectance properties. The interpreter thus makes an informed decision whether the spectral change was caused by a mortality event, or is the result of clouds, noise, vegetation phenology or other ephemeral changes not related to structural changes of the forest canopy. Hence, the final measurement recorded for each plot and year was the presence or absence of a canopy mortality event. The manual interpretation of temporal-spectral profiles is a well-established method (Cohen *et al*. 2010) that yields more precise estimates of annual canopy mortality rates than automated algorithms (Cohen *et al*. 2017). The approach has been successfully applied across many forest ecosystems globally (Pflugmacher *et al*. 2012; Hermosilla *et al*. 2015; Potapov *et al*. 2015; Cohen *et al*. 2016; Senf *et al*. 2018). Yet, the final call for each plot and year remains a human decision and is thus prone to measurement errors similar to measurements taken in the field. While most mortality events will result in well-identifiable spectral changes that are easy to detect (see Figure S2 for an example), it might be particularly challenging to detect low severity mortality events that have small spectral changes in relation to the noise inherent to satellite time series. While we cannot rule out the omission of such low severity mortality events, we aimed to make their detection as consistent as possible: For each plot we ran an automatic change detection algorithm (Kennedy *et al*. 2010) and compared the outcome to our human interpretation. If there was an inconsistency between the human and automatic interpretation, the most knowledgeable interpreter revisited the plot to check whether the initial interpretation was correct. If an error was observed, we corrected the initial interpretation. While this procedure is not able to rule out all potential measurement errors, it guarantees a high degree of consistency in the assessment, as plots that were particularly hard to interpret were collectively interpreted by the same interpreter.

From the number of plots experiencing canopy mortality we estimated annual canopy mortality rates using a Bayesian partially pooled binomial model with a logit link function (Senf *et al*. 2018) implemented in Stan (Carpenter *et al*. 2017). In essence, the model estimates the annual rate of plots experiencing a mortality event over the total number of plots per country using repeated binary trials (hence the binomial likelihood function). The model further includes a linear regression term with year as predictor, explicitly modeling the fractional change (through the logit link) in the mean canopy mortality rate over time (i.e., the temporal trend in canopy mortality). The partial pooling applied to our model assumes each year’s mortality rate to emerge from the same underlying distribution, which is beneficial when estimating rates in repeated measurements as it shrinks individual estimates towards the mean (Gelman *et al*. 2014). This shrinkage towards the mean will prevent large outliers (e.g., a year with very high mortality due to a storm or an extraordinary fire season) to dominate the trend, and also reduces the impact of potential outliers related to interpretation errors on the overall result. We demonstrate the robustness of our model to omitted disturbances in a sensitivity analysis presented in Figures S3 and S4. To derive estimates at the country, regional and continental scale, we first modeled annual canopy mortality rates and trends at the country level, subsequently aggregating to regions (Central-, Eastern-, Northern-, South-Eastern, Southern-, and Western-Europe; Fig. S1) and the continental level using forest cover as weight (i.e., accounting for the stratified sampling design; Table S1).

### Simulating future forest demography

We used neutral landscape models (NLM) to assess the impact of different future mortality trajectories on forest demography. An NLM constitutes a minimal model with regard to the underlying assumptions about the processes that drive landscapes. Yet they have proven to be powerful tools for assessing critical thresholds in landscape properties (e.g., the occurrence of old forests; Synes *et al*. 2016). As our aim was to determine how trends in forest canopy mortality could potentially affect forest demography, NLMs provide a robust and parsimonious approach. We built NLMs with two different landscape configurations (random and clumped forest distribution) for each country using the NLRM package (Sciaini *et al*. 2018) in R. The grain of the simulation was set to 1 ha (i.e., 100 ×100 m), which is close to the median patch size of natural mortality in temperate Europe (Senf and Seidl 2018). The extent of the simulations was fixed at 100 km^2^ (i.e., 10 × 10 km), resulting in 10,000 potentially forested cells (hereafter referred to as ‘stands’) simulated per country. The proportion of forested stands was set to the average forest proportion of each country. We mapped forest age classes for the year 2015 to stands using reconstructed age class distributions from Vilén *et al*. (2012). For six countries, age class data were not available (Bosnia and Herzegovina, Greece, Serbia, Montenegro, Northern Macedonia, and Moldova), which limited the simulation-based analyses to 29 countries.

After initializing the landscapes, we iteratively updated the age of each stand on a yearly basis (i.e., aging them by one year) over the period 2019-2050. The annual proportion of pixels experiencing mortality was drawn each year from the Bayesian partially pooled model calibrated from observed canopy mortalities (described in the previous section). We hence used an empirical mortality model calibrated from observed data to simulate mortality rates for each country. To allocate mortality in space we ranked each stands’ likelihood of being affected by mortality based on four alternative age-based mortality functions (Figure S5). Individual tree mortality is U-shaped over stand age, with high mortality risk in young trees (competition for resources, self-thinning) and old trees (hydraulic limitations, disturbances like wind and insects, timber harvesting). As tree mortality from competition between individual trees does not leave a strong signature in the forest canopy and is thus unlikely to be detected in our satellite-based approach to identifying canopy mortality at a grain of 30 × 30 m, we focused on the latter (i.e., the mortality of old trees) in our simulations. We hence simulated a monotonically increasing canopy mortality probability with age in our NLMs (Figure S5). Furthermore, we assumed no recovery failure in our simulations, meaning all stands regrew after a mortality event.

We used NLMs to study three scenarios: First, a stabilization of canopy mortality rates at the mean value observed for the period 1985 to 2018; second, a stabilization of canopy mortality rates at values observed for 2018; and third, a further increase in canopy mortality at the country-specific trends observed for the past 34 years (see Figure S6). In total, we ran eight alternative NLM configurations per country (two landscape configurations × four alternative mortality functions) for three scenarios, which were replicated 15 times for each of the 29 countries, resulting in the analysis of 10,440 NLMs in total. From those NLMs, we derived the median age of forests within each country for the year 2050 as an indicator of changes in forest demography in response to the different mortality scenarios.

## Results and discussion

### Trends in canopy mortality rates

Canopy mortality increased by 1.50 ± 0.28 % per year across Europe (Figure 1). The average canopy mortality rate in the late 20^th^ century (1985 to 1999) was 0.79 ± 0.04 % yr^-1^ (i.e., a forest area of 1.7 Mill. ha affected by canopy mortality each year) and increased to 0.99 ± 0.04 % yr^-1^ (i.e., 2.1 Mill. ha) in the early 21^st^ century (2000 to 2018). Mortality increased at an accelerating pace throughout the observation period, with the highest canopy mortality rate of the past 34 years observed in 2018 (1.14 ± 0.16 % yr^-1^).

**Fig. 1.**
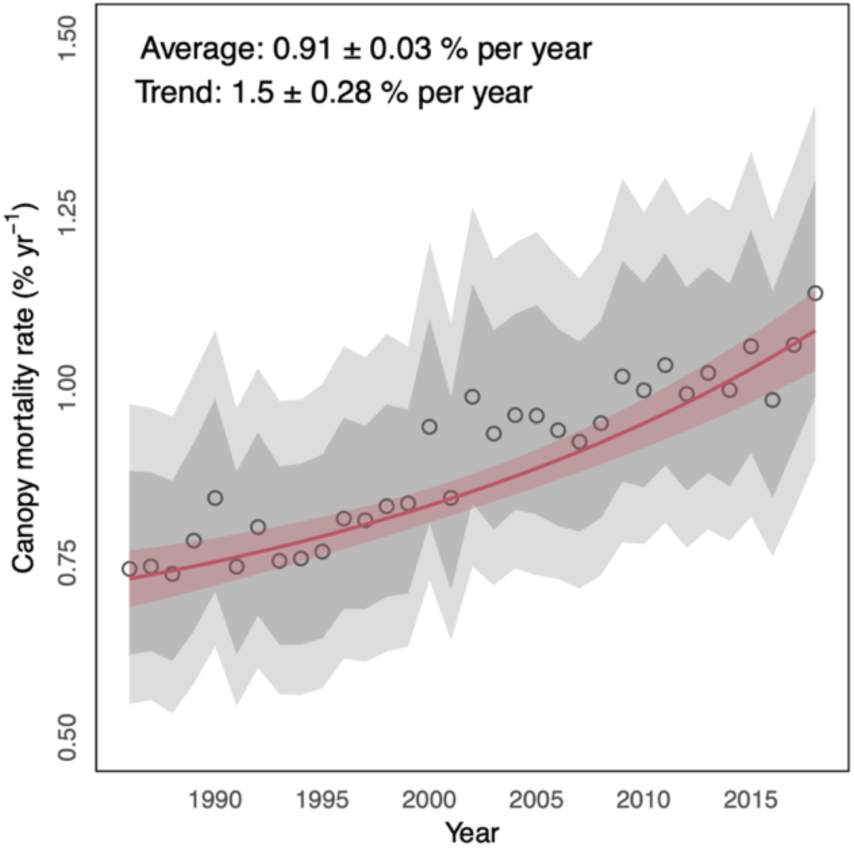
Canopy mortality rates and trends in Europe’s forests. Black circles indicate the mean continental-scale canopy mortality, with grey ribbons indicating the uncertainty range (darker grey = standard deviation; lighter grey = 90 % credible interval). The red line gives the mean temporal trend and its standard deviation.

Changes in canopy mortality varied substantially between countries and regions (Figure 2). Out of the 35 countries analyzed, 28 had a positive trend in canopy mortality (Figure 2; Table S2 and Figure S7). Trends were strongest in Central and Eastern Europe, where canopy mortality increased on average by 55 % and 78 % from the late 20^th^ to the early 21^st^ century. Weaker but nonetheless positive trends were found for Western and Northern Europe, where canopy mortality increased on average by 39 % and 19 % over the same period. No evidence for changes in canopy mortality could be found for South-Western Europe, and evidence for increasing canopy mortality in South-Eastern Europe was weak (Figure 2).

**Fig. 2.**
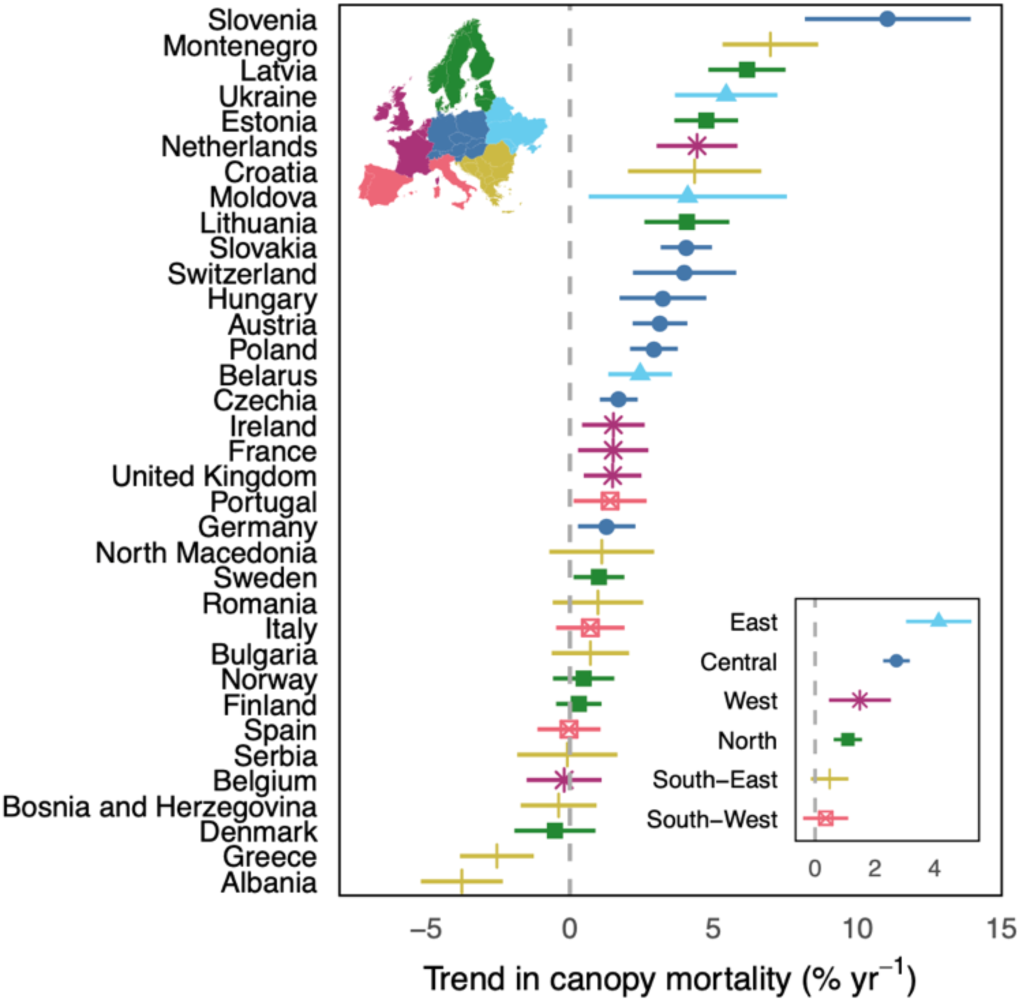
Trends in canopy mortality at the country and regional level. Dots indicate the mean trend in canopy mortality 1985-2018, error-bars give the standard deviation. Colors indicate the assignment of countries to different regions (see inserts).

Drivers of increasing canopy mortality are manifold, interactive, and vary locally. While an in-depth analyses of these local drivers is beyond the scope of this study, we here discuss several coinciding continental-scale developments that are likely to contribute to the overall increase in canopy mortality observed for Europe: First, large parts of Europe’s forests have accumulated biomass in recent decades (Ciais *et al*. 2008), and many countries have started utilizing their growing timber resources more actively by increasing annual fellings (European Environmental Agency 2016). The notion that timber harvest is an important factor contributing to increasing canopy mortality is also supported by a high correlation of our estimates with data from wood harvesting statistics (Figure S8 and S9). Human resource use is thus likely to be a major driver of current trends in canopy mortality identified for Europe’s forests. Second, the collapse of the Soviet Union has led to large-scale transformations of economic and political systems in parts of Europe, resulting in a pronounced increase in both regular harvests and illegal logging (Kuemmerle *et al*. 2007). The particularly strong increase in canopy mortality rates observed for European countries of the former Soviet Union (Figure S4) can in part be explained by those historical legacies. Third, many forests across Europe have seen episodes of large-scale storm events and severe bark beetle outbreaks in recent decades (Seidl *et al*. 2014), which likely contribute to the particularly strong trends observed for Central and Eastern Europe. Hence, increased natural disturbances – a result of both structural legacies and climate change (Seidl *et al*. 2011) – can thus be considered an additional important driver of increasing canopy mortality trends. Increased tree mortality caused by drought has further been reported for Europe recently (Allen *et al*. 2010; Carnicer *et al*. 2011), and the particularly high mortality rates observed for 2018 might be a consequence of the recent drought affecting large parts of Europe (Buras *et al*. 2020; Schuldt *et al*. 2020). In this context it is interesting to note, however, that we did not find strong evidence for increasing canopy mortality in Southern Europe, despite the notion that warm and dry areas are particularly prone to drought-induced forest dieback (Allen *et al*. 2010). In the past, drought-induced forest dieback in Mediterranean systems might have still been too dispersed to trigger substantial changes in canopy mortality as observed from satellite imagery (Hartmann *et al*. 2018). Likewise, while increasing forest fire activity is expected across the Mediterranean (Moriondo *et al*. 2006) and the actual number of fires has increased in the past, the annual area burned has decreased in recent decades (San-Miguel-Ayanz *et al*. 2013; see also Figure S10). Despite this overall decrease in burned area across southern Europe, the years 2017 and 2018 were both characterized by unprecedentedly strong fire seasons (San-Miguel-Ayanz *et al*. 2018, 2019), which likely further contributed to the particularly high mortality rates in Europe’s forests in 2017 and 2018. Finally, we note that only 2.6 % of the observed mortality events led to a change in land use from forest to non-forest. That is, land use change is of relatively minor importance for forest canopy mortality in Europe.

### Impacts of increasing forest canopy mortality on forest demography

We found widely varying demographic trajectories in the three scenarios of future forest mortality in Europe (Figure S11). A stabilization of canopy mortality at the level observed in the past (1985-2018) would not result in drastic demographic changes (Figure 3). This scenario in fact increases the proportion of countries with considerable amounts of old forests (i.e., median forest age of 60 years or older) compared to the current situation. The aging trend currently observed in Europe’s forests thus outweighs the effects of tree mortality on demography in this scenario. Stabilizing canopy mortality rates at the level observed for 2018, however, stops the aging trend of Europe’s forests (Figure 3), leading to an increase of countries with young forests (median age younger than 30 years). A continued increase in canopy mortality at rates as observed for the past 34 years would lead to considerable shifts in the age structure of Europe’s forests (Figure 3). For instance, while the forests of only 14 % of European countries have a median age younger than 30 years today, this number increases to 53 ± 3 % in 2050 if observed canopy mortality trends will continue. A continued increase in canopy mortality would thus inverse the current aging trend of Europe’s forests.

**Fig. 3.**
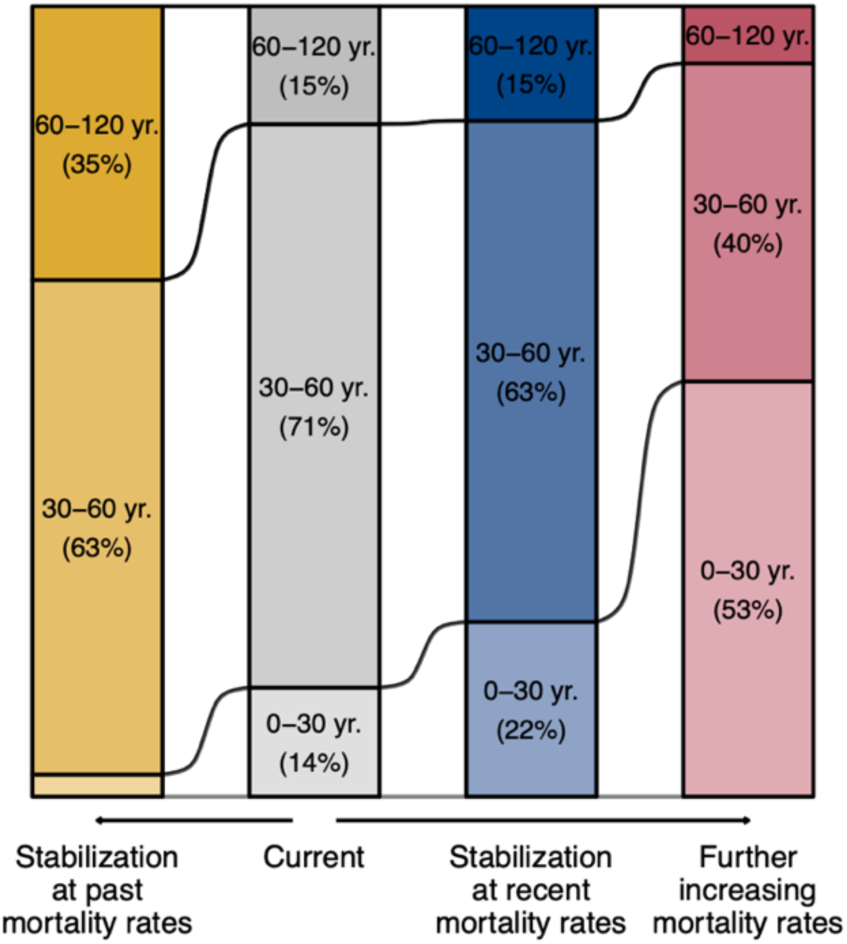
Effects of changing canopy mortality rates on the median age of Europe’s forests in 2050. The distribution of median ages of the forests of 29 European countries was simulated for the year 2050 under three scenarios: (1) Stabilization at canopy mortality rates observed between 1985 and 2018 (*Stabilization at past rates*); (2) Stabilization at canopy mortality rates observed for the year 2018 (*Stabilization at recent rates*); (3) Increasing canopy mortality rates extrapolating trends observed between 1985 and 2018 for each country (*Further increasing mortality rates*) (see Figure S6 for details on the three scenarios). The current condition shows the distribution of median ages in 2015. Percentage values show the distribution of countries in different bins of median age. We here show the averages over all landscape configurations and mortality functions, see Figure S12 for the results of individual landscape configurations and mortality functions. See Figure S11 for a detailed representation of individual runs of the NLMs for each country.

Changes in forest demography resulting from increasing canopy mortality have the potential to challenge the integrity of Europe’s forests in at least three important ways. First, a shift towards young forests could result in considerable bottlenecks for the regeneration of forests due to decreasing seed availability and increased distance to seed sources because of a decreasing share of mature forests on the landscape (Anderson-Teixeira *et al*. 2013). Second, the prevalence of old forests will be substantially reduced under a continued increase in canopy mortality. Such a shift towards younger forests simultaneously reduces the diversity in age classes on the landscape, resulting in a biological homogenization of forest habitats (van der Plas *et al*. 2016). As both the prevalence of old forests and the diversity in developmental stages are important indicators of biodiversity (Hilmers *et al*. 2018), a future increase in canopy mortality can have widespread negative consequences for forest biodiversity. Third, increasing canopy mortality reduces the residence time of carbon in forest ecosystems (Yu *et al*. 2019; Pugh *et al*. 2019), with negative impacts on the total carbon stored in forests (Körner 2017). Likewise, old forests are hotspots of forest carbon storage and act as long-term carbon sinks (Zhou *et al*. 2006; Luyssaert *et al*. 2008). The loss of old forests under increasing canopy mortality rates would thus reduce the carbon storage potential of Europe’s forests. A continued increase in canopy mortality might hence offset potential C gains from accelerated tree growth under climate change (Yu *et al*. 2019).

## Conclusion and management implications

Here we show that canopy mortality rates have increased consistently throughout the past 34 years in Europe, and that a further increase has the potential to substantially alter Europe’s forest demography. It is of paramount importance for forest policy and management to counteract the ongoing trends in forest mortality and implement strategies to safeguard the integrity of Europe’s forest ecosystems. This could be achieved via: (i) increasing the resistance and resilience of Europe’s forests to natural disturbances by, e.g., counteracting biotic homogenization; (ii) conserving existing and creating future old-growth forests by establishing areas that are exempt from timber extraction, especially in places where the risk of natural disturbances is low; (iii) accounting for natural disturbances in long-term forest planning; and (iv) considering demographic constraints in managing forests for a bio-based economy. We conclude that developing strategies to address the increasing canopy mortality should be a key priority of forest policy and management in Europe.

## Acknowledgments

We thank Alice Cosatti, Barbara Öllerer, Benjamin Stadler, Laura Steinbach and Lukas Weikl for their help with the interpretation of the satellite data. We further thank Dirk Pflugmacher and Yang Zhiqiang for their help with implementing Google Earth Engine code. C. Senf acknowledges funding from the Austrian Science Fund (FWF) Lise-Meitner Program (Nr. M2652). R. Seidl and J. Sebald acknowledge funding through FWF START grant Y895-B25. We are grateful for helpful comments from two anonymous reviewers.

## Author contributions

C. Senf and R. Seidl acquired the funding for the research project. C. Senf and R. Seidl conceptualized the research project. C. Senf and J. Sebald administrated the research project and curated the data acquisition. C. Senf conducted the formal data analysis. C. Senf wrote the original draft, with review and editing from J. Sebald and R. Seidl.

## Data and code availability

All data and code used in the analysis will be made available via a public repository upon publication of the article. Requests for material during review should be directed to C. Senf.

