## Supplementary Materials for "Increases in canopy mortality and their impact on the demographic structure of Europe’s forests"

**Table S1:** Countries included in the analysis and realized sample sizes per country.

| Country | Land area ($km^2$) | Forest area ($km^2$) | Realized sample size |
|---|---|---|---|
| Albania | 28,786 | 7,716 | 415 |
| Austria | 83,988 | 39,600 | 1828 |
| Belarus | 207,575 | 80,334 | 442 |
| Belgium | 30,587 | 6,834 | 334 |
| Bosnia and Herzegovina | 51,030 | 25,599 | 393 |
| Bulgaria | 110,953 | 36,250 | 418 |
| Croatia | 57,017 | 24,901 | 433 |
| Czechia | 78,842 | 26,000 | 1401 |
| Denmark | 43,501 | 6,120 | 401 |
| Estonia | 45,405 | 23,066 | 377 |
| Finland | 338,434 | 233,320 | 450 |
| France | 549,006 | 246,640 | 413 |
| Germany | 357,454 | 114,190 | 1227 |
| Greece | 124,885 | 37,600 | 412 |
| Hungary | 93,001 | 20,990 | 349 |
| Ireland | 70,243 | 7,540 | 298 |
| Italy | 300,887 | 106,736 | 363 |
| Latvia | 64,549 | 28,807 | 389 |
| Lithuania | 64,941 | 21,223 | 416 |
| Moldova | 33,847 | 3,290 | 302 |
| Montenegro | 13,764 | 6,252 | 403 |
| Netherlands | 35,162 | 3,650 | 322 |
| Norway | 311,654 | 121,120 | 394 |
| Poland | 311,759 | 90,000 | 1312 |
| Portugal | 88,700 | 31,820 | 287 |
| Romania | 238,289 | 69,610 | 436 |
| Serbia | 88,372 | 27,200 | 334 |
| Slovakia | 49,036 | 20,006 | 1820 |
| Slovenia | 20,221 | 12,574 | 412 |
| Spain | 498,518 | 184,180 | 326 |
| Sweden | 450,040 | 280,730 | 439 |
| Switzerland | 41,239 | 12,540 | 1128 |
| North Macedonia | 25,438 | 10,285 | 439 |
| Ukraine | 597,120 | 105,000 | 451 |
| United Kingdom | 245,164 | 28,650 | 332 |
| **Total** | **5,749,424** | **2,100,373** | **19,896** |

**Table S2:** Average canopy mortality rates and fractional changes for the period 1985-2018 per region and country. Reported are the mean plus/minus standard deviation of the posterior distribution.

| Region and country | Average canopy mortality rate (% per year) | | | Fractional change in canopy mortality rate (% per year) |
|---|---|---|---|---|
| | 1985-2018 | 1985-1999 | 2000-2018 | |
| Central Europe | 0.82±0.03 | 0.62±0.04 | 0.96±0.05 | 2.72±0.45 |
| Austria | 1.22±0.08 | 0.86±0.08 | 1.48±0.13 | 3.13±0.95 |
| Czechia | 1.20±0.08 | 1.01±0.10 | 1.35±0.11 | 1.70±0.66 |
| Germany | 0.85±0.07 | 0.73±0.09 | 0.94±0.11 | 1.28±1.00 |
| Hungary | 0.70±0.11 | 0.46±0.13 | 0.87±0.16 | 3.24±1.51 |
| Poland | 0.56±0.05 | 0.41±0.06 | 0.66±0.07 | 2.92±0.83 |
| Slovakia | 1.05±0.08 | 0.63±0.07 | 1.35±0.12 | 4.05±0.89 |
| Slovenia | 0.34±0.07 | 0.08±0.05 | 0.54±0.12 | 11.05±2.88 |
| Switzerland | 0.66±0.09 | 0.39±0.07 | 0.86±0.16 | 3.99±1.80 |
| Easter Europe | 0.58±0.06 | 0.40±0.07 | 0.71±0.09 | 4.14±1.10 |
| Belarus | 0.78±0.10 | 0.61±0.13 | 0.90±0.14 | 2.44±1.11 |
| Moldova | 0.29±0.08 | 0.15±0.08 | 0.39±0.12 | 4.10±3.45 |
| Ukraine | 0.43±0.07 | 0.24±0.08 | 0.57±0.11 | 5.43±1.78 |
| Northern Europe | 1.07±0.06 | 0.97±0.09 | 1.15±0.09 | 1.10±0.48 |
| Denmark | 0.81±0.11 | 0.79±0.16 | 0.82±0.14 | -0.52±1.41 |
| Estonia | 0.89±0.12 | 0.53±0.13 | 1.15±0.17 | 4.74±1.11 |
| Finland | 1.25±0.13 | 1.22±0.18 | 1.27±0.16 | 0.31±0.79 |
| Latvia | 1.32±0.14 | 0.66±0.15 | 1.80±0.22 | 6.16±1.34 |
| Lithuania | 0.60±0.09 | 0.39±0.11 | 0.74±0.13 | 4.07±1.47 |
| Norway | 0.81±0.11 | 0.77±0.16 | 0.83±0.15 | 0.48±1.06 |
| Sweden | 1.07±0.12 | 0.96±0.16 | 1.15±0.16 | 1.01±0.89 |
| South-Eastern Europe | 0.48±0.03 | 0.46±0.05 | 0.49±0.04 | 0.49±0.63 |
| Albania | 0.70±0.10 | 0.96±0.17 | 0.52±0.11 | -3.76±1.43 |
| Bosnia and Herzegovina | 0.48±0.08 | 0.49±0.12 | 0.47±0.11 | -0.39±1.32 |
| Bulgaria | 0.72±0.10 | 0.65±0.14 | 0.77±0.14 | 0.71±1.35 |

| | | | | |
|---|---|---|---|---|
| Croatia | 0.21±0.05 | 0.13±0.06 | 0.28±0.08 | 4.34±2.32 |
| Greece | 0.63±0.10 | 0.76±0.16 | 0.54±0.12 | -2.54±1.29 |
| Montenegro | 1.01±0.12 | 0.40±0.11 | 1.46±0.19 | 6.97±1.66 |
| North-Macedonia | 0.48±0.08 | 0.44±0.11 | 0.52±0.11 | 1.11±1.82 |
| Romania | 0.34±0.07 | 0.31±0.10 | 0.37±0.09 | 0.97±1.57 |
| Serbia | 0.34±0.08 | 0.34±0.11 | 0.33±0.10 | -0.09±1.74 |
| South-Western Europe | 1.09±0.09 | 1.04±0.13 | 1.12±0.12 | 0.35±0.76 |
| Italy | 0.76±0.11 | 0.70±0.16 | 0.8±0.15 | 0.71±1.19 |
| Portugal | 2.63±0.23 | 2.27±0.33 | 2.89±0.31 | 1.40±1.27 |
| Spain | 1.01±0.13 | 1.03±0.20 | 1.00±0.17 | -0.03±1.10 |
| Western Europe | 0.97±0.10 | 0.79±0.14 | 1.10±0.14 | 1.50±1.04 |
| Belgium | 0.76±0.11 | 0.76±0.17 | 0.76±0.15 | -0.20±1.30 |
| France | 0.96±0.12 | 0.77±0.16 | 1.10±0.17 | 1.50±1.23 |
| Ireland | 1.01±0.14 | 0.87±0.19 | 1.11±0.19 | 1.51±1.09 |
| Netherlands | 0.95±0.14 | 0.57±0.15 | 1.23±0.20 | 4.42±1.41 |
| United Kingdom | 1.06±0.13 | 0.92±0.18 | 1.17±0.18 | 1.48±1.01 |

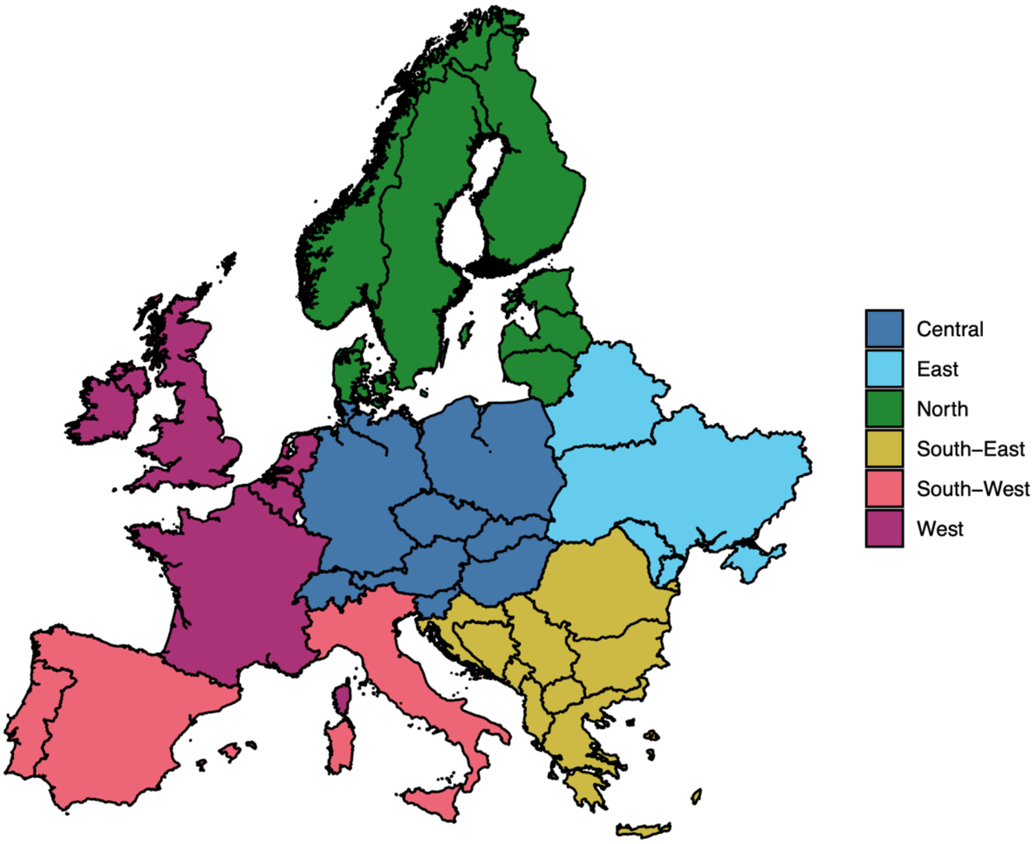

**Fig. S1:** Countries and regions included in the analysis.

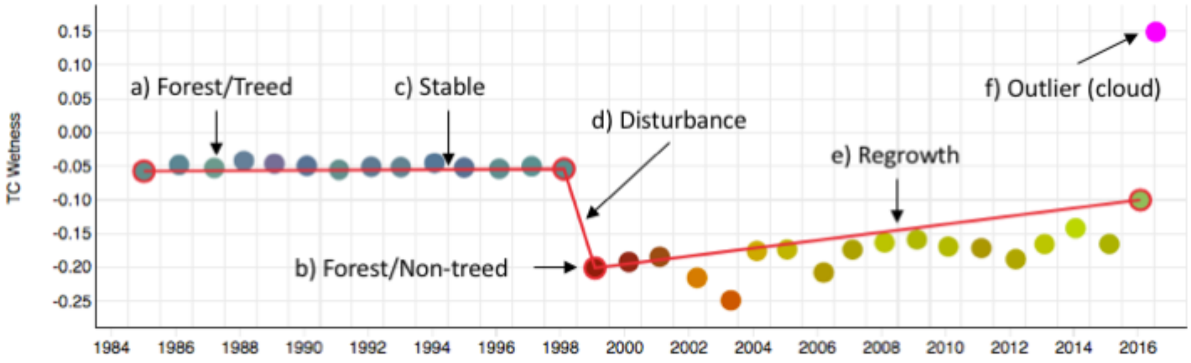

**Figure S2:** Example of a spectral-temporal profile. The dots indicate annual Landsat observations in Tasseled Cap space (i.e., an ordination of the six spectral bands of Landsat into three independent components of brightness, greenness and wetness that can be visualized in RGB-space). The y-axis value is the Tasseled Cap wetness value, which is highly sensitive to structural changes in the tree canopy (but other indices can be selected by the interpreter). The time series was segmented into three linear segments by the interpreter, presenting stable conditions, declining conditions (mortality caused by a disturbance), and increasing conditions (regrowth). Each segment is defined by two vertices, at which the land use and land cover are recorded. Outliers (clouds) are easily spotted by their very bright color (see year 2017). From the segmented time series, the occurrence of a mortality event can be estimates for each year, which is later used to estimate annual mortality rates.

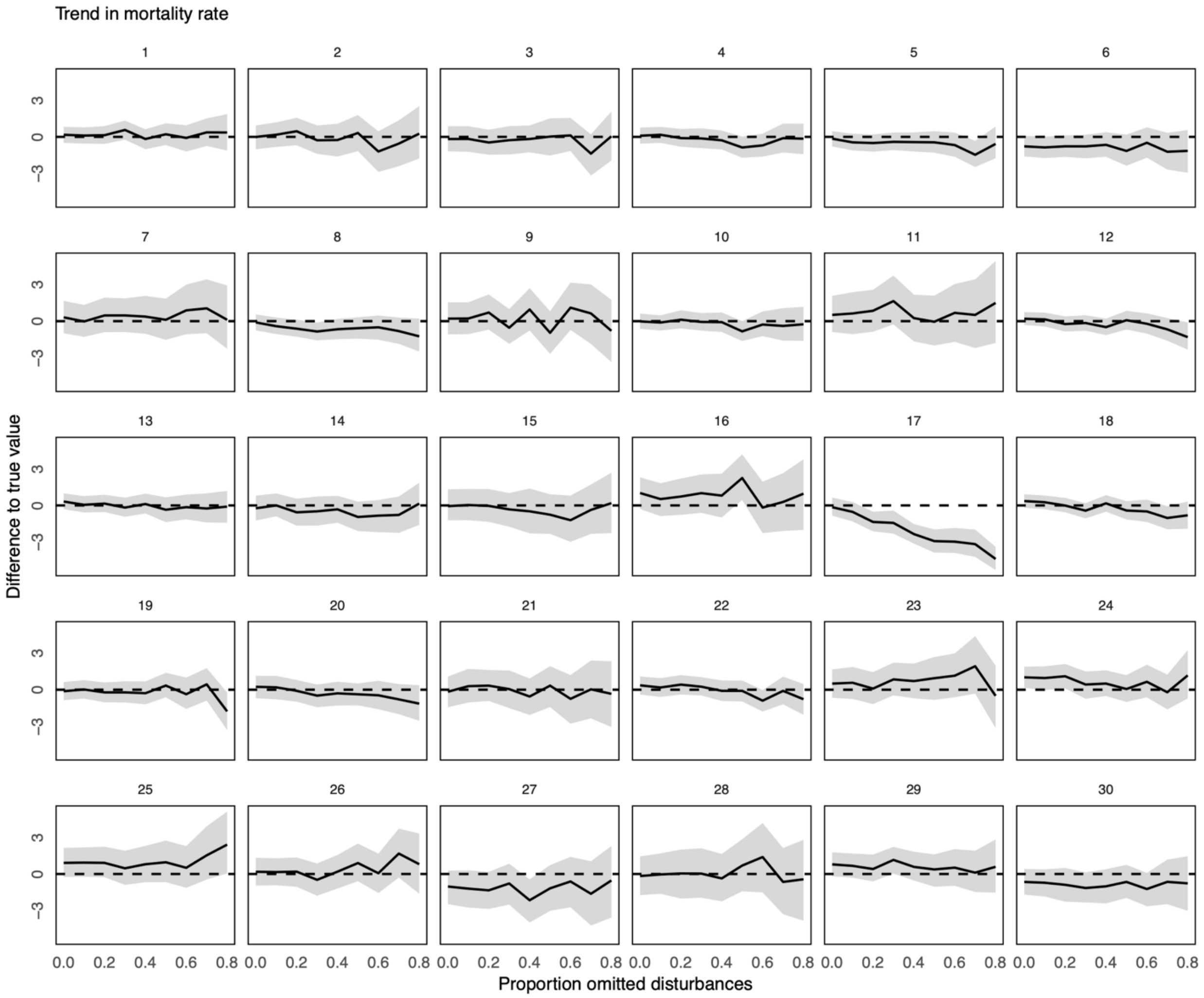

**Figure S3:** Sensitivity analysis of trend estimates. For the sensitivity analysis, we simulated data from the Bayesian partially-pooled model using known parameters. The true mortality rate rate was simulated using a half-normal distribution with a mean of 0.01 and a standard deviation of 0.05. The true variation among years was simulated using a half-normal distribution with a mean of 0.1 and a standard deviation of 0.05. The true trend was simulated using a normal distribution with a mean of 0.04 and a standard deviation of 0.05. We used a Bayesian partially-pooled model to estimate the true parameters, but sequently increased the number of omitted mortality events (i.e., simulating that the interpreter makes an error in identifying a canopy mortality event). We repeated the simulation 30 times (panels 1 through 30) and report the difference between the estimate and the true value (y-axis). Black lines indicate the median of the posterior and grey ribbons indicate the 95 % credible interval. Dashed horizontal lines indicate no deviation from the true value. From this sensitivity analysis it is evident that our trend estimates are highly robust against omitted mortality events during interpretation. That is, there was only one instance where omitting mortality events systematically biased the estimate (run number 17). For all other instances, omitting mortality events only decreased the precision of the estimate but did not bias the estimate itself.

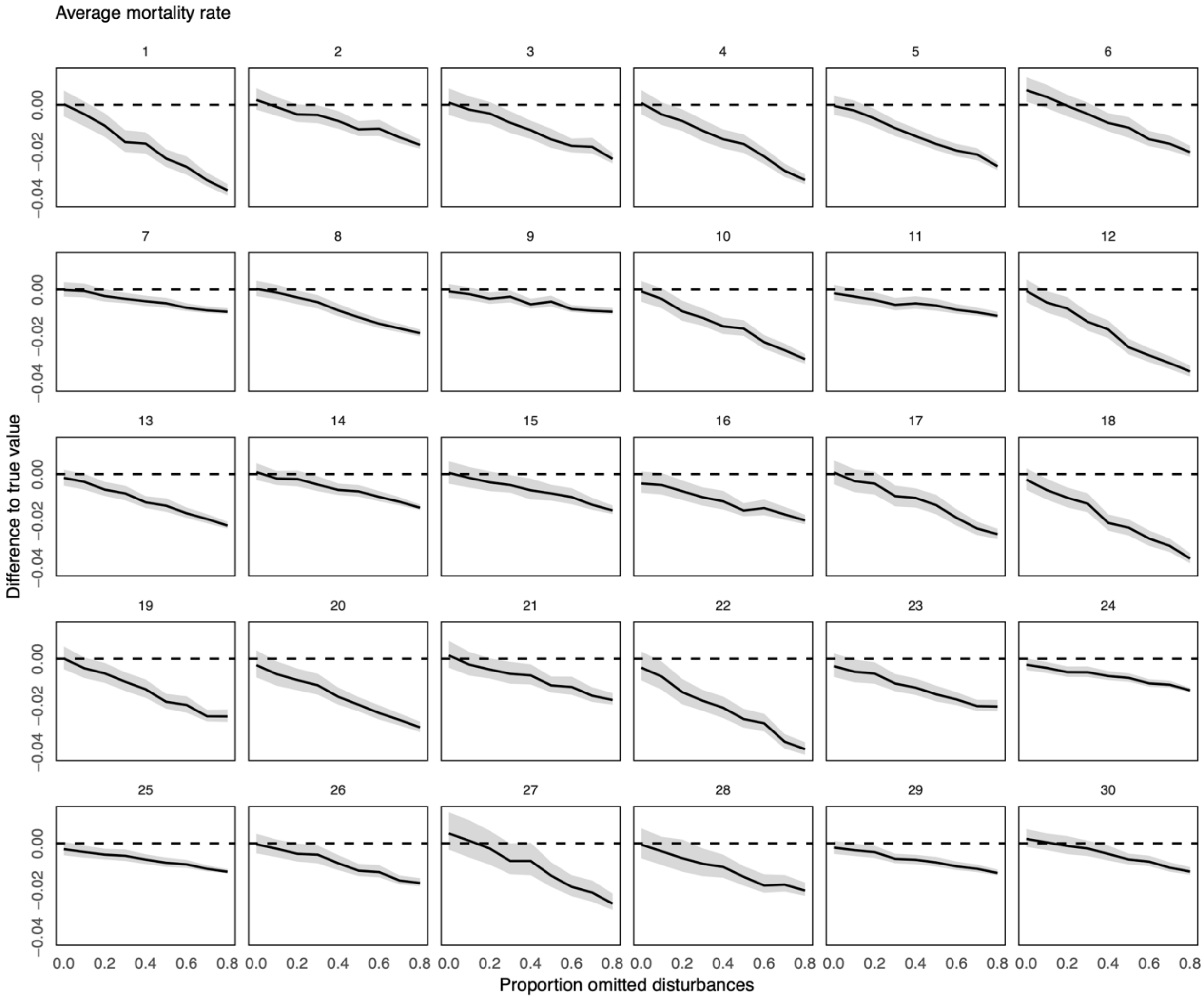

**Figure S4:** Sensitivity analysis of average canopy mortality rates. For this sensitivity analysis, we simulated data from the Bayesian partially-pooled model using known parameters. The true mortality rate was simulated using a half-normal distribution with a mean of 0.01 and a standard deviation of 0.05. The true variation among years was simulated using a half-normal distribution with a mean of 0.1 and a standard deviation of 0.05. The true trend was simulated using a normal distribution with a mean of 0.04 and a standard deviation of 0.05. We used a Bayesian partially-pooled model to estimate the true parameters, but sequently increased the number of omitted mortality events (i.e., simulating that the interpreter makes an error in identifying a canopy mortality event). We repeated the simulation 30 times (panels 1 through 30) and report the difference between the estimate and the true value (y-axis). Black lines indicate the median of the posterior and grey ribbons indicate the 95 % credible interval. Dashed horizontal lines indicate no deviation from the true value. In contrast to the trend estimates (Figure S3) the estimates of the average mortality rate were affected by omitted mortality events during interpretation. As expected, the average mortality rate estimated from the data decreased with increasing omission of mortality events. However, significant bias only occurred when more than 20 % of the true mortality events were omitted. We conclude that our estimates of average mortality rates can be seen as conservative (i.e., if they are biased by interpreter errors, they likely underestimate the true rates).

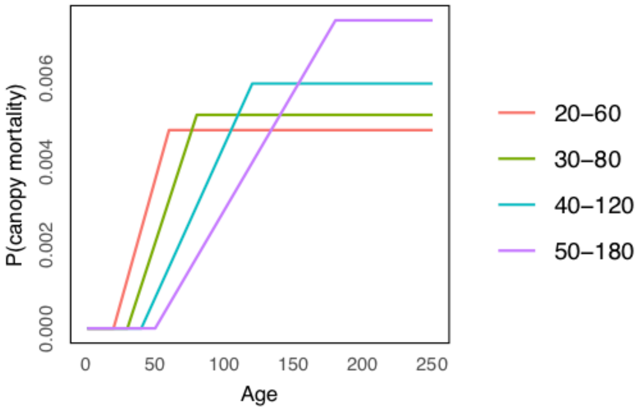

**Fig. S5:** Alternative age-based mortality functions considered in the neutral landscape models.

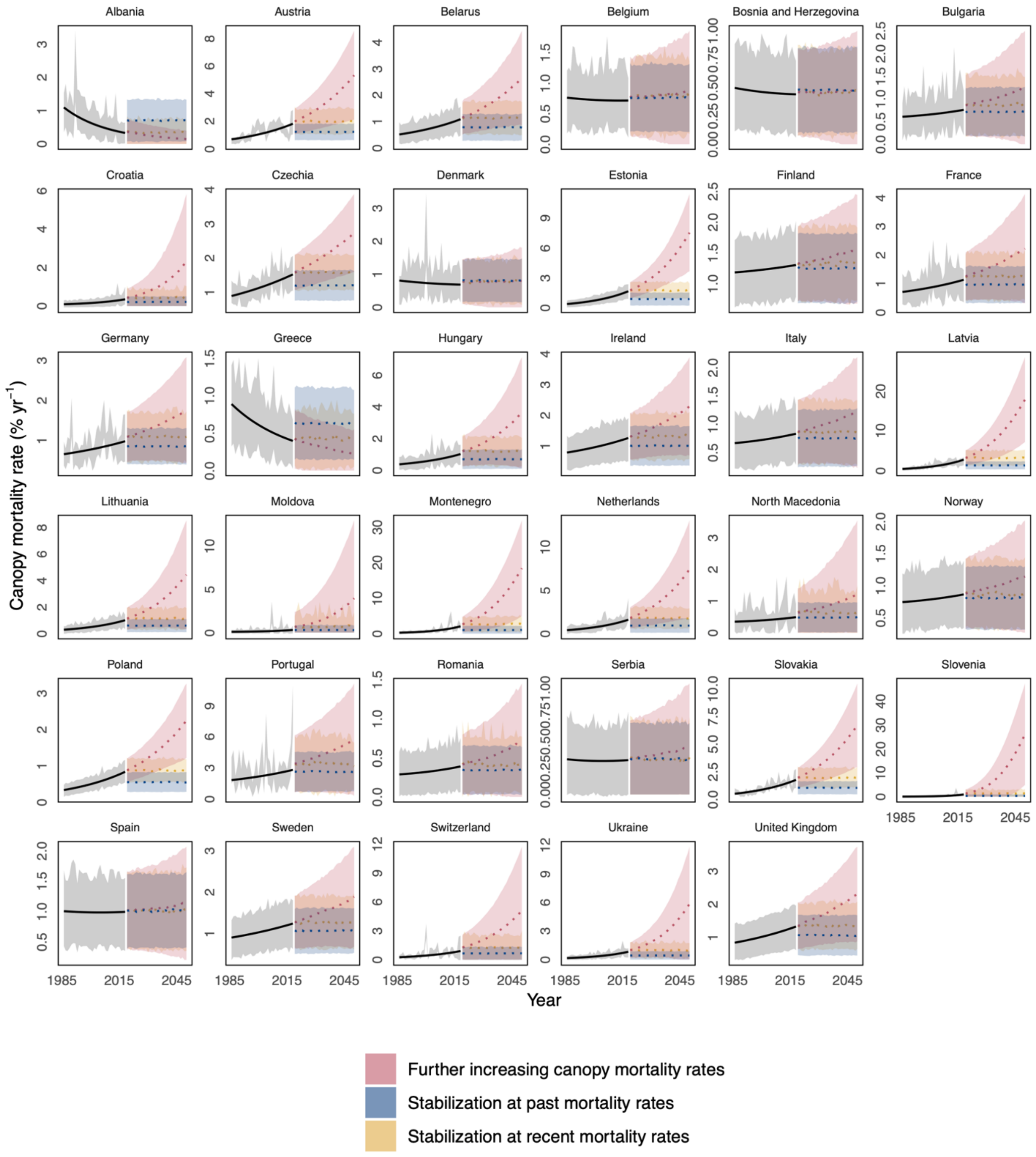

**Fig. S6:** Scenarios of future canopy mortality rates: (1) Further increasing canopy mortality rates, continuing trends as observed for the period 1985-2018 (red); (2) Stabilization at past mortality rates (1986-2018; blue); (3) Stabilization at recent rates (2018; green). Uncertainty bands show the standard deviation of the posterior, while lines show the mean.

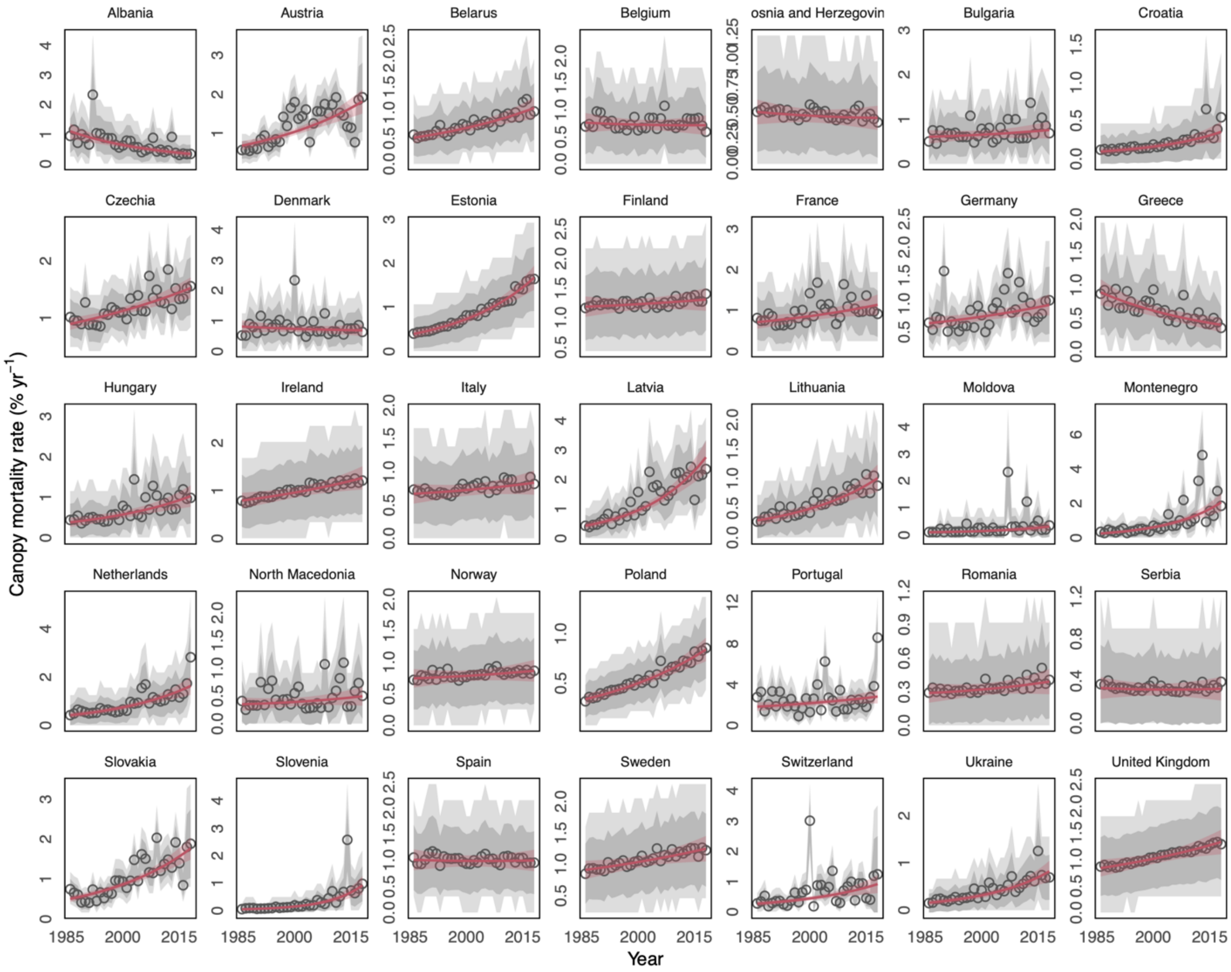

**Fig. S7:** Annual canopy mortality rates estimates for 35 European countries. Black circles indicate the average annual canopy mortality, with grey ribbons indicating the uncertainty range (darker grey = standard deviation; lighter grey = 90 % credible interval). The red line gives the average temporal trend with ribbons indicating its standard deviation. Note that y-axes are scaled differently in each panel.

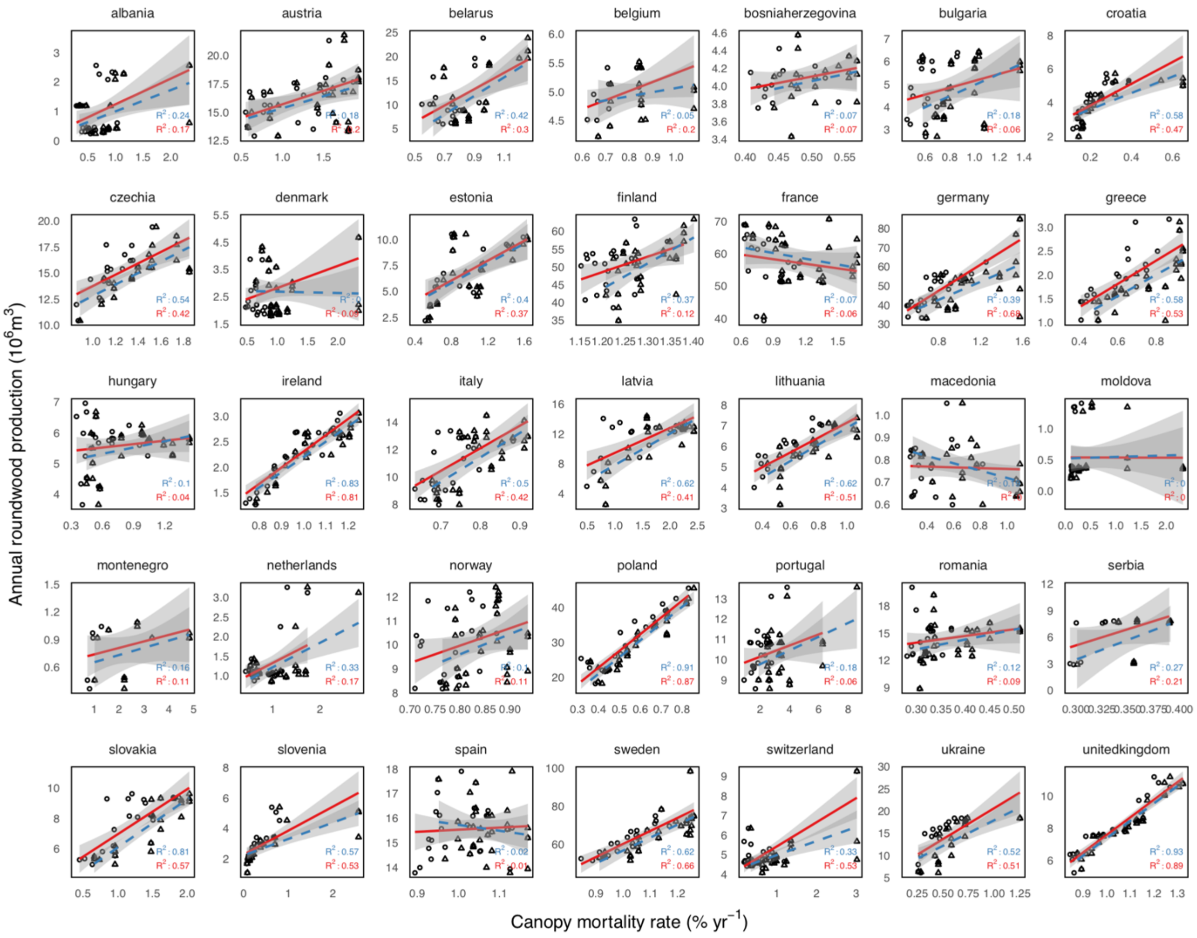

**Fig. S8:** Correlation between annual canopy mortality rates and annual roundwood extraction rates as reported by FAOSTAT (http://www.fao.org/faostat/en/#home). Circles and solid red lines represent the year-to-year correlation, whereas triangles and dashed blue lines indicate correlations between annual FAOTSTAT data and a 3-year moving maximum of canopy mortality rates, accounting for potential differences in the attribution of canopy mortality events to individual years.

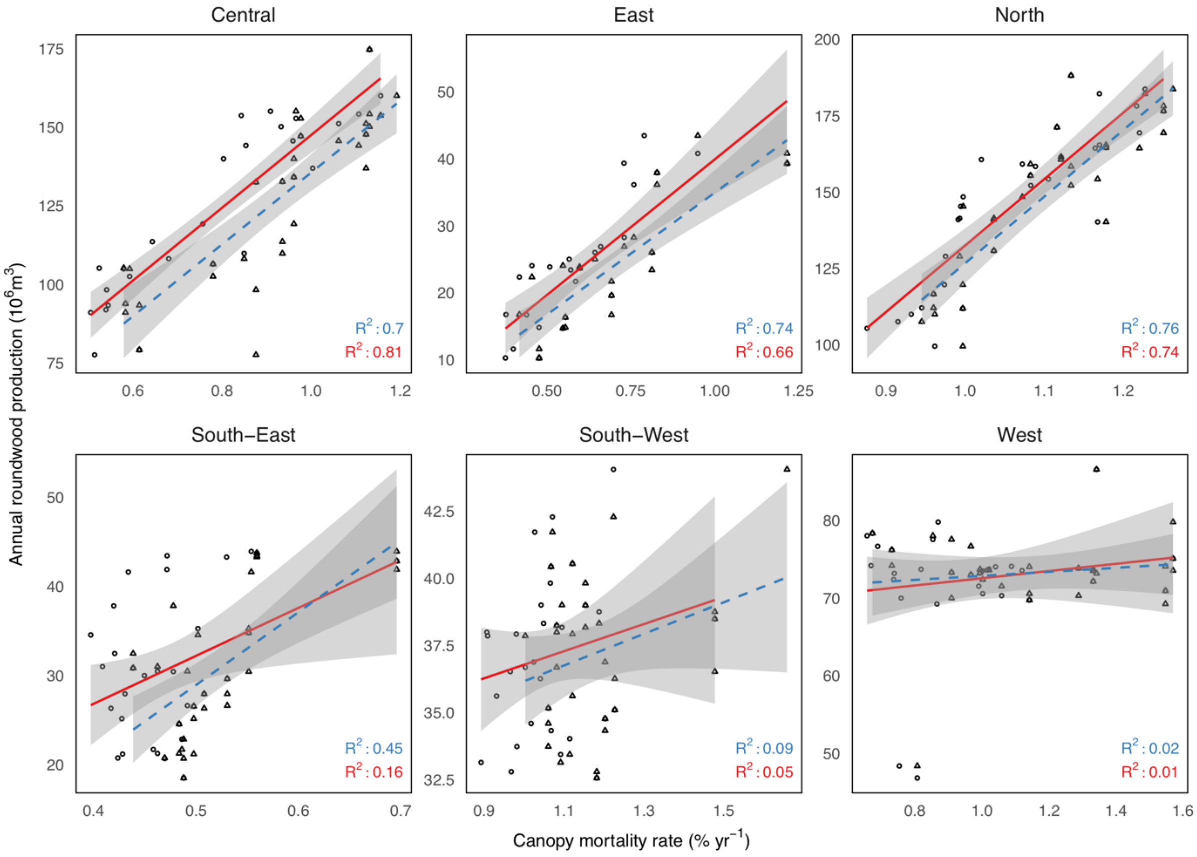

**Fig. S9:** Same as Fig. S8 but aggregated to regional level.

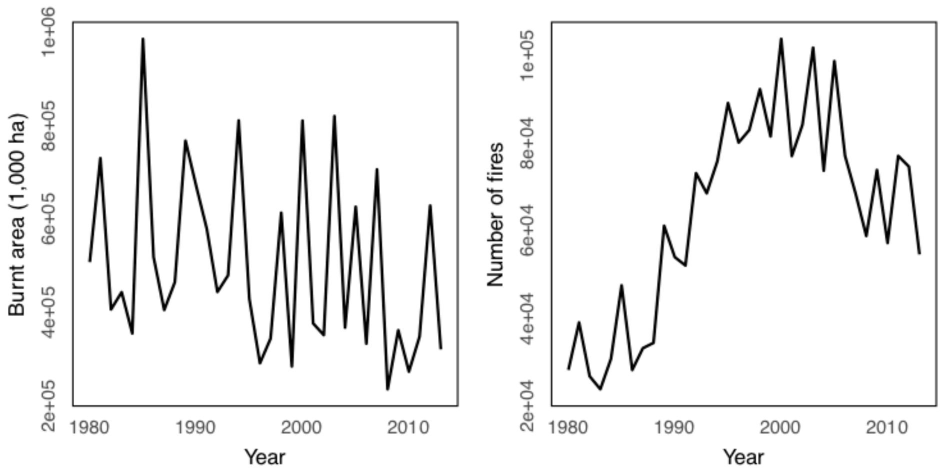

**Fig. S10:** Total burnt area and number of fires across Southern Europe (France, Greece, Italy, Portugal and Spain), as reported by the European Environmental Agency (https://www.eea.europa.eu/data-and-maps/indicators/forest-fire-danger-2/assessment).

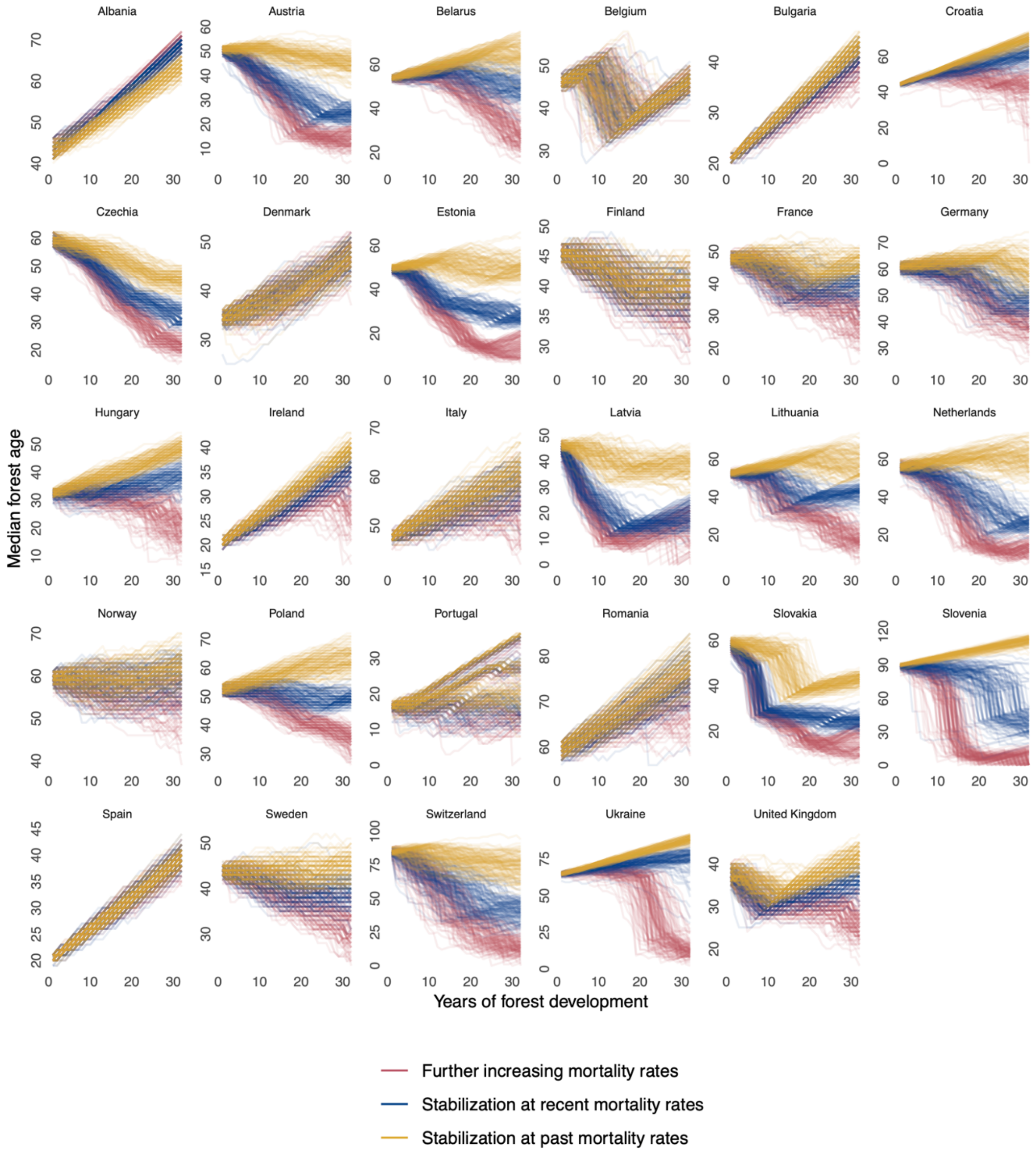

**Fig. S11:** Individual neutral landscape model (NLM) runs for all countries where data was available. Each line presents the median age simulated over 32 years (2018 to 2050) by one NLM run. Colors indicate the three future mortality scenarios (see Fig. S3 for scenarios).

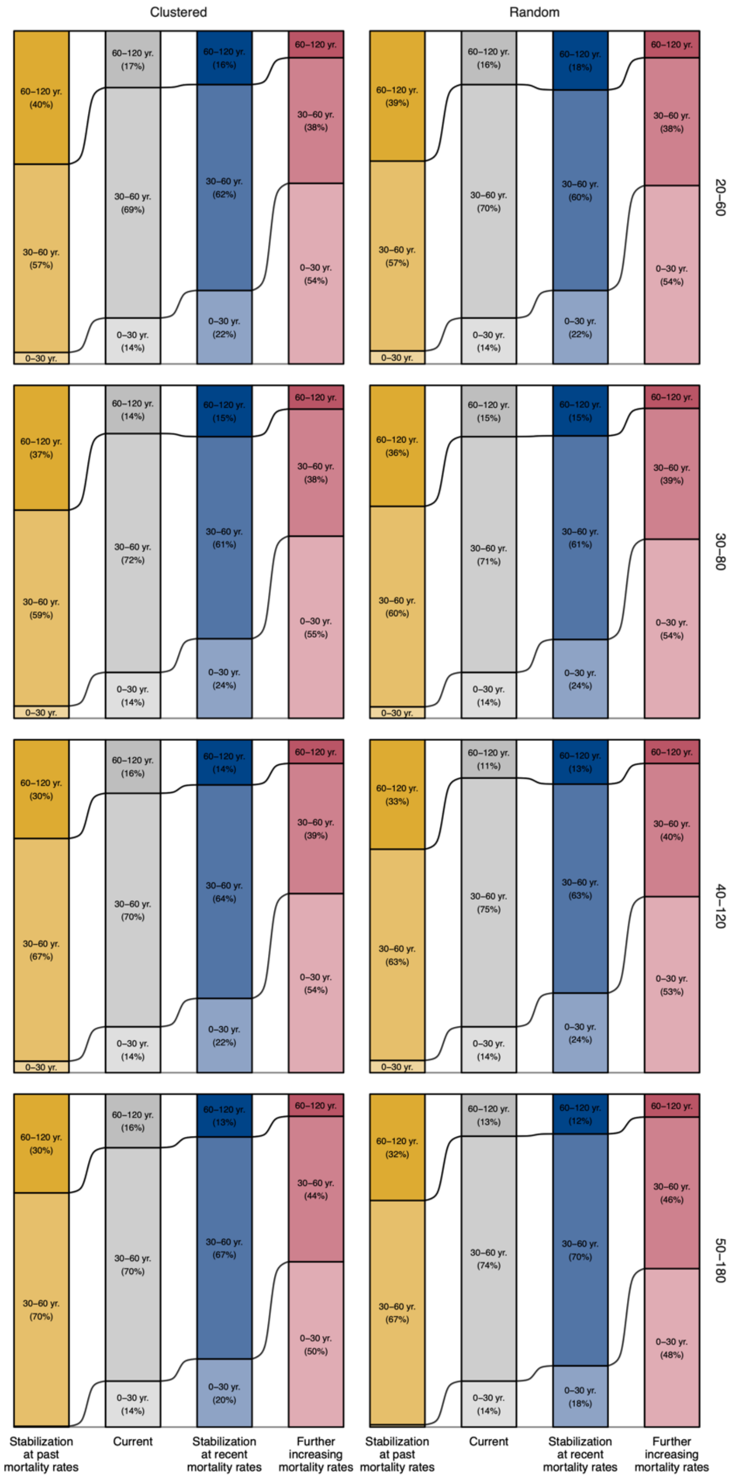

**Fig. S12:** Changes in median age for different initial configurations of neutral landscape models. Columns show different landscape initializations (random versus clumped) and rows are different mortality functions (see Fig. S2).

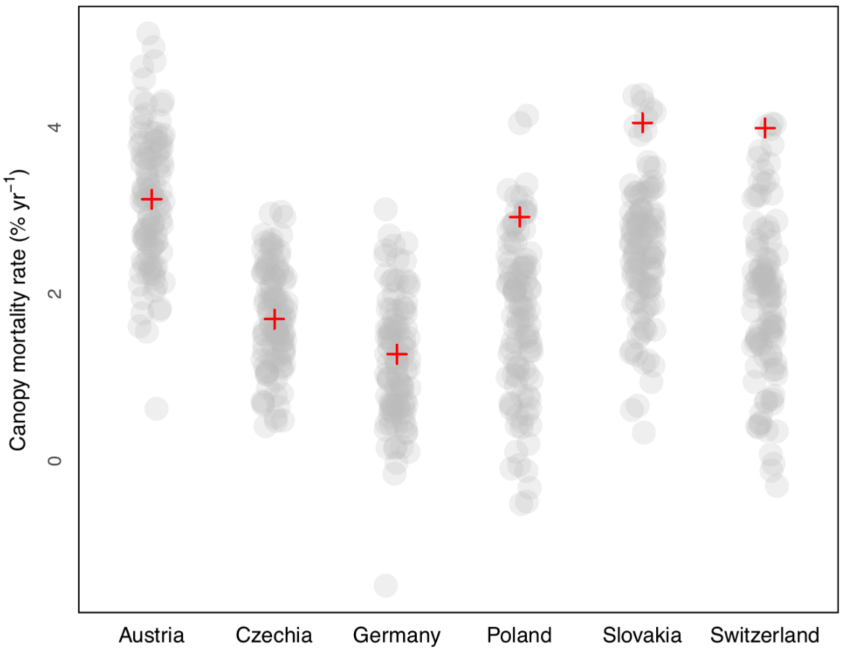

**Fig. S13:** Effects of down-sampling data for six countries in Central Europe where larger sample sizes were available to estimate canopy mortality trends. Grey dots in the background show mortality trend estimates from 100 model runs with reduced sample size (n = 500). The red crosses are mortality trend estimates with the full sample (see Tale S1 for sample sizes).
